# Structure of Ycf1p reveals the transmembrane domain TMD0 and the regulatory R region of ABCC transporters

**DOI:** 10.1101/2021.01.29.428621

**Authors:** Sarah C. Bickers, Samir Benlekbir, John L. Rubinstein, Voula Kanelis

## Abstract

ATP binding cassette (ABC) proteins typically function in active transport of solutes across membranes. The ABC core structure is comprised of two transmembrane domains (TMD1 and TMD2) and two cytosolic nucleotide binding domains (NBD1 and NBD2). Some members of the C-subfamily of ABC (ABCC) proteins, including human multidrug resistance proteins (MRPs), also possess an N-terminal transmembrane domain (TMD0) that contains five transmembrane α-helices and is connected to the ABC core by the L0 linker. While TMD0 was resolved in SUR1, the atypical ABCC protein that is part of the hetero-octameric ATP-sensitive K+ channel, little is known about the structure of TMD0 in monomeric ABC transporters. Here, we present the structure of yeast cadmium factor 1 protein (Ycf1p), a homologue of human MRP1, determined by electron cryomicroscopy (cryo-EM). Comparison of Ycf1p, SUR1, and a structure of MRP1 that showed TMD0 at low resolution demonstrates that TMD0 can adopt different orientations relative to the ABC core, including a 145° rotation between Ycf1p and SUR1. The cryo-EM map also reveals that segments of the regulatory (R) region, which links NBD1 to TMD2 and was poorly resolved in earlier ABCC structures, interacts with the L0 linker, NBD1, and TMD2. These interactions, combined with fluorescence quenching experiments of isolated NBD1 with and without the R region, suggests how post-translational modifications of the R region modulate ABC protein activity. Mapping known mutations from MRP2 and MRP6 onto the Ycf1p structure explains how mutations involving TMD0 and the R region of these proteins lead to disease.

**Statement of Significance:** The Ycf1p structure provides an atomic model for the TMD0 domain of ABCC transporters and for two segments of the regulatory (R) region that links NBD1 to TMD2. The orientation of TMD0 in Ycf1p differs from that seen in SUR1, the regulatory ABCC protein in KATP channels, demonstrating flexibility in TMD0/ABC core contacts. The structure suggests how post-translational modifications of the R region modulate ABC protein activity and provides a mechanistic understanding of several diseases that occur due to mutation of human homologues of Ycf1p.

## Introduction

ATP binding cassette (ABC) proteins are a large family of membrane proteins found in all kingdoms of life (1, 2). ABC proteins have a core structure comprised of two transmembrane domains (TMD1 and TMD2) and two cytosolic nucleotide binding domains (NBD1 and NBD2) (Fig. 1A, Fig. S1A) (3–5). Through ATP binding and hydrolysis at the NBDs, ABC proteins actively transport solutes across cell membranes, regulate activities of other proteins, or function as channels (1, 2). Thus, ABC proteins are involved in many biological processes, including lipid homeostasis, cellular metal trafficking, and antigen peptide transport. Mutations in human ABC proteins cause diseases such as Tangier disease, adenoleukodystrophy, cystic fibrosis, Dubin-Johnson syndrome, and pseudoxanthoma elasticum (1, 2). Further, the export of a wide range of cancer chemotherapeutics, antibiotics, and anti-fungal drugs by ABC transporters confers multi-drug resistance to tumor cells, bacteria, and fungal pathogens, respectively (1, 2, 6, 7).

**Fig. 1.**
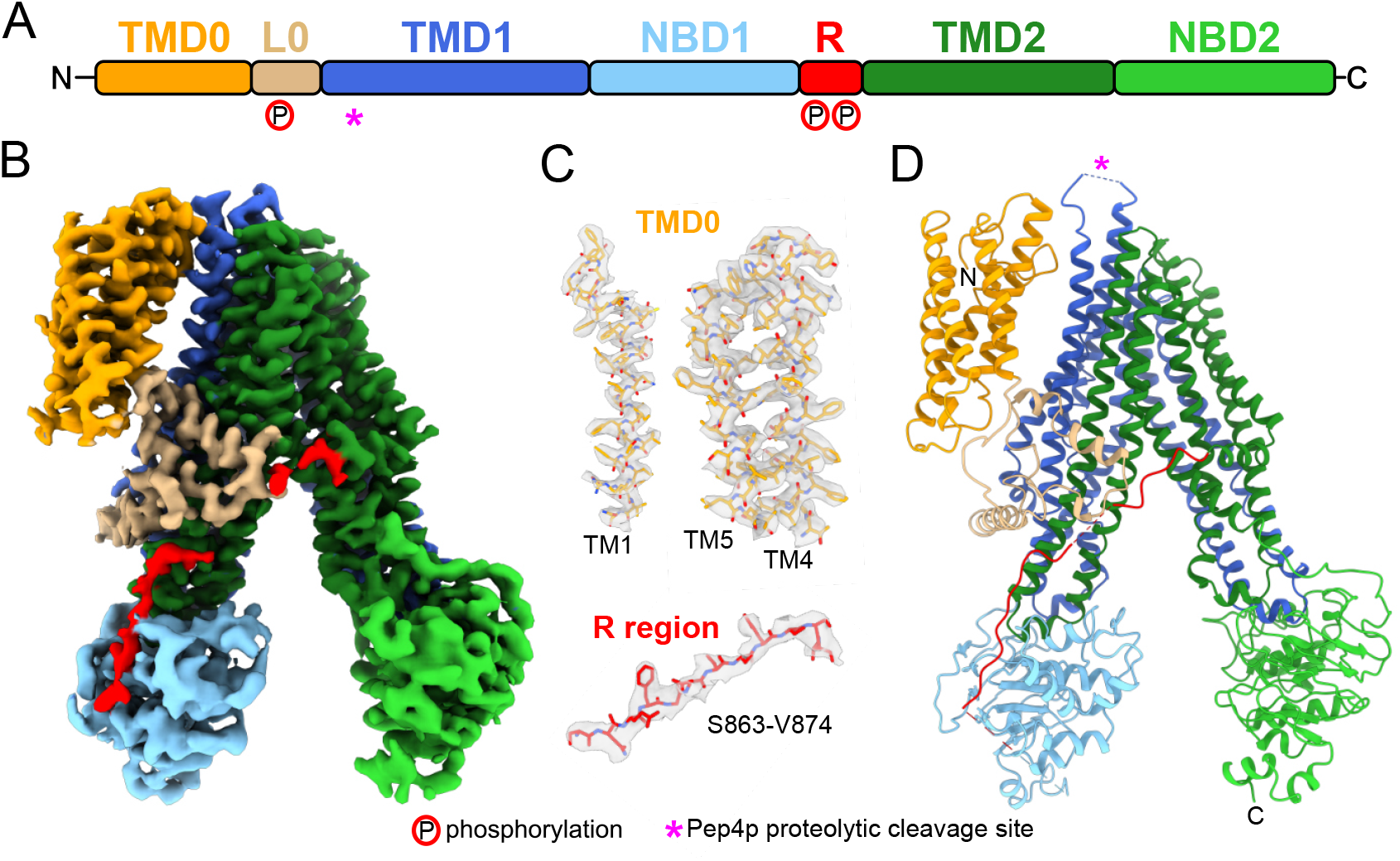
Ycf1p Structure. (A) Ycf1p domain arrangement. TMD, transmembrane domain; L0, L0 linker; NBD, nucleotide binding domain; R, regulatory region. The Pep4p proteolytic digestion site within lumenal loop 6 of TMD1 is denoted by a pink “ * ”. Phosphorylation sites in the L0 linker (S251) and R region (S908 and T911) are depicted with a “P” circled in red. (B) Cryo-EM density of Ycf1p with domains coloured as in panel A. (C) Example of the atomic model for individual TM helices in TMD0 and the R region fit into the corresponding map densities. (D) Schematic ribbon diagram of Ycf1p coloured as in panels A and B and with the proteolytic digestion site denoted by a pink “ * ”.

Human ABC proteins are divided into seven subfamilies (A-G), based in part on the sequence of their NBDs and TMDs in the core ABC structure (1, 2). The C subfamily is the most diverse and includes the cystic fibrosis transmembrane conductance regulator (CFTR), the sulphonylurea receptors (SURs) that form regulatory subunits in ATP-sensitive K+ (KATP) channels, and the multidrug resistance proteins (MRPs). In addition to the ABC core, ABCC proteins contain an N-terminal extension that is either comprised of an additional transmembrane domain (TMD0) and L0 linker (Fig. 1A, *orange* and *tan*, respectively, and Fig. S1A) or just an L0 tail (5, 8). A TMD0, but not L0 linker, is also found in some ABCB proteins (3, 5). These N-terminal extensions are involved in trafficking, endosomal recycling, protein interactions, and/or regulation of ABC activity (9–18). The existence of disease-causing mutations in TMD0 and L0 linker of different ABCC proteins (8, 13, 18) indicates that TMD0 and the L0 linker play important roles in protein function when present.

High-resolution structural information for TMD0 is available only for the atypical ABCC protein SUR1 (19, 20), which is part of the large hetero-octameric ATP-sensitive K+ (KATP) channel complex. In contrast, structures of monomeric ABC transporters showed only low-resolution density for TMD0 that was insufficient for building a full atomic model or lacked density for the domain altogether (14, 21–24). The vacuolar ABCC protein yeast cadmium factor 1 (Ycf1p) from *Saccharomyces cerevisiae* is a close homologue of human MRPs and a model for ABCC proteins that function as monomers. Ycf1p transports glutathione-conjugated heavy metals, such as Cd^2+^, from the cytosol into the vacuole, detoxifying the cell (25, 26). Human MRP1 can rescue Cd^2+^ transport activity in a *YCF1* deletion strain (27).

Like other ABCC proteins, Ycf1p contains a relatively long and mostly disordered linker that connects NBD1 and TMD2 (25, 28, 29) (Fig. 1A, Fig. S1A). This linker contains stimulatory phosphorylation sites, similar to the phospho-regulatory (R) region in the ABCC protein CFTR (30–32). However, how the R region interacts with the ABCC core, and how its phosphorylation modulates protein function remain poorly understood for most ABCC proteins. Structural studies of Ycf1p presented here reveal how TMD0 and the R region exert their regulatory functions in MRP-like ABCC proteins.

## Results and Discussion

### Overall Ycf1p structure

To isolate Ycf1p for structural analysis by electron cryo-microscopy (cryo-EM), sequence encoding a 3×FLAG affinity tag was integrated into the yeast chromosomal DNA at the 3’ end of the *YCF1* gene. Following growth and harvest of the resulting yeast strain, membranes were solubilized with detergent and Ycf1p was isolated by affinity chromatography. Although not required for Ycf1p function, proteolytic digestion of Ycf1p at an insertion in loop 6 (Fig. S1A) can affect the substrate specificity of the protein (33). However, the yeast strain used for protein purification lacked the Pep4p protease, and consequently Ycf1p was isolated without proteolytic digestion, as confirmed by SDS-PAGE (Fig. S2A). The Ycf1p preparation showed a specific ATPase activity of 32.2 ± 3.9 nmol P*i*/min/mg protein (± s.d., n=6 replicates, with three replicates from each of two protein batches from independent yeast cultures, Fig. S2B).

Cryo-EM of the sample led to a map of the protein with an overall resolution of 3.2 Å (Fig. 1B and C, Fig. S3A to D). Local resolution of the protein ranges from ~2.7 to ~6.7 Å. The TMDs (TMD0, TMD1, and TMD2) and L0 linker are at ~2.7 to ~3.8 Å resolution, allowing construction of atomic models for these regions (Fig. 1C and Fig. S3E, S4, S5). The NBDs are at ~4.7 to ~6.7 Å resolution and were interpreted by fitting of homology models of Ycf1p NBD1 and NBD2 (Fig. S3E, S5). The quality of the map also enabled construction of an atomic model for two segments of the R region from residues S863 to R875 and H928 to K935 (Fig. 1C, Fig. S5). The atomic model for Ycf1p comprises 94 % of its 1515 residues (L12-N324, H360-L852, S863-R875, H928-V1512), making it the most complete model of a monomeric ABCC protein to our knowledge.

In the structure, the ABC core of Ycf1p adopts the inward-facing conformation (Fig. 1B and D, Fig. S1B) seen for eukaryotic ABCB, ABCC, and ABCD transporters, as well as some bacterial exporters, when bound nucleotides are absent (5). TMD1 and TMD2 each contain six transmembrane (TM) segments and are domain-swapped to form two TM bundles that contain residues from both halves of the ABC core (Fig. S1B). TM bundle 1 is comprised of TM helices 6, 7, 8, 11, 15, and 16, while TM bundle 2 is formed by TM helices 9, 10, 12, 13, 14, and 17. The TM domain swapping results in coupling helices 1 and 4 (CH1 and CH4) contacting NBD1 and CH2 and CH3 contacting NBD2. The cytosolic NBDs are asymmetrically separated, similar to other ABCC transporters in the inward conformation (8). The inward conformation of Ycf1p also exposes a number of residues likely involved in glutathione binding (Fig. S6) (22). In addition, several bound lipid-molecules are found throughout the transmembrane domains (Fig. S7A), including one at the interface between TMD0 and TM bundle 1, which is modelled as phosphatidyl ethanolamine (Fig. S7B, C), a common lipid in yeast vacuolar membranes (34, 35).

### High-resolution structure of TMD0 and the L0 linker

The hallmark of ABCC transporters is their N-terminal extension, which is either comprised of TMD0 and L0 linker or an L0 tail alone (1, 29) and which has various functions, depending on the ABCC protein (9, 10, 13, 15–17). For example, the N-terminal extensions of MRP2 (9) and CFTR (16) are involved in localization of the proteins to the plasma membrane, and trafficking of these ABCC proteins is regulated through interactions of the N-terminal extension with vesicle fusion machinery (16, 17). TMD0 and L0 in SUR1 (13) and MRP1 (10) mediate protein-protein interactions, while the MRP1 L0 linker can affect transport activity (15). In Ycf1p, TMD0 is required for vacuolar localization and is thought to form interactions with regulatory protein partners (36, 37). The Ycf1p L0 linker is essential for the protein’s role in cadmium resistance (36).

The structure of the Ycf1p TMD0 shows the domain’s five α-helices and N-terminal tail (1, 29) (Fig. 1 and 2, *orange*). The Ycf1p L0 linker adopts a lasso structure that consists of a membrane-embedded region (R207-S251), an amphipathic helix (S252-Q269) that packs against the elbow helix of TMD1, and an L0 linker/ABC core connecting segment (K270-S274) (Fig. 1 and 2A, *tan*) (5, 8).

**Fig. 2.**
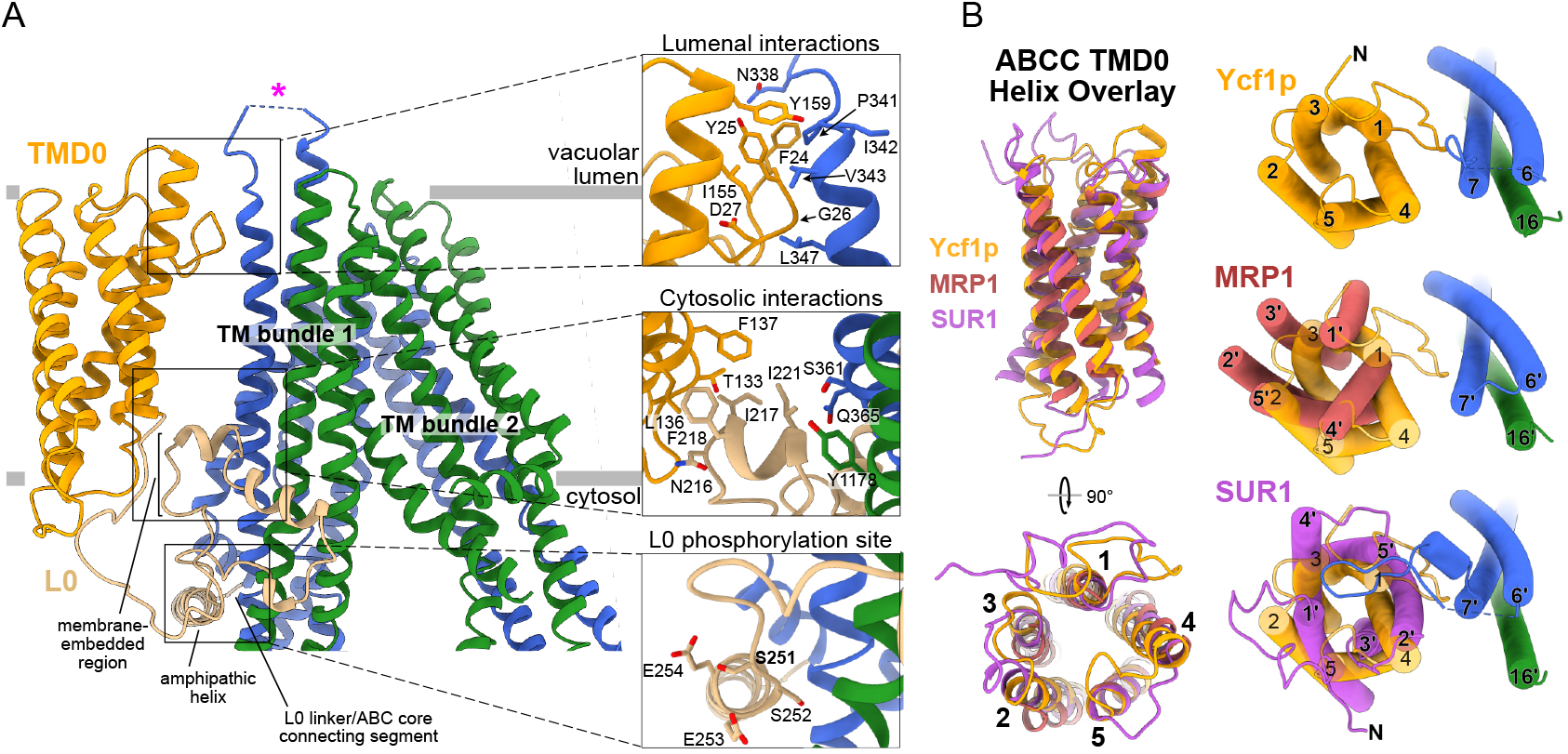
TMD0 and L0 interactions in Ycf1p and other ABCC proteins. (A) A Ycf1p ribbon diagram coloured as in Fig. 1 (left). Close-up views of the luminal interaction of TMD0 and TM bundle 1 (top right) and cytoplasmic interaction of TMD0, the L0 linker, and TM bundle 1 (middle right) are shown with interacting side chains (<4 Å apart) rendered as sticks (right). A close-up view of the L0 linker phosphorylation site with side chains rendered as stick is also shown (bottom right). (B) Overlay of TMD0 from Ycf1p, MRP1, and SUR1 (left) shows that the structure of TMD0 is conserved among these different ABCC proteins. Overlay of the TMD0 (right) from Ycf1p (shown in orange), MRP1 (shown in red), and SUR1 (shown in purple) after alignment of the ABC core demonstrates that the orientation of TMD0 differs in ABCC proteins. TM helix numbers in MRP1 (middle right) and SUR1 (bottom right) are denoted with a “ ′ ” to distinguish them from Ycf1p TM helices.

The lumenal and cytoplasmic sides of TMD0 contact TM bundle 1 and the L0 linker, respectively, to tether TMD0 to the rest of Ycf1p. On the lumenal side of the membrane (Fig. 2A, top right), residues in the N-terminal tail and TM4 in TMD0 contact residues in TM6 and loop 6 of TMD1. Loop 6, which connects TM6 and TM7 in TMD1, contains the Pep4p protease digestion site (33). While most residues in loop 6 are not resolved in the structure, it is possible that this loop interacts transiently with TMD0 and that these interactions change upon proteolytic digestion. In support of this hypothesis, loop 6 in SUR1 contacts TMD0 in some KATP channel structures (18, 38), but is not observed in other KATP channel structures (19, 20, 39), likely due to flexibility. On the cytoplasmic side of the membrane, there is a network of interactions formed by TM4 and TM5 in TMD0, the membrane-embedded portion of the L0 lasso structure, the L0 amphipathic helix, and residues in TM6, TM7, TM15, and TM16 in TM bundle 1 (Fig. 2A, centre and bottom right). Notably, TMD0 contacts two regulatory elements in Ycf1p – loop 6 that can be proteolytically cleaved and the L0 linker that contains an inhibitory phosphorylation site – suggesting that the TMD0 interactions may change with different transporter states.

The lasso structure of the L0 linker has implications for Ycf1p activity. The S251 inhibitory phosphorylation site in the Ycf1p L0 linker is 70 % buried within the protein (Fig. 2A, bottom right), making it inaccessible to kinases in this conformation. Further, the side chains of two Glu residues C-terminal to S251 that are thought to be involved in recognition by casein kinase II (CKII) (40), the *in vivo* kinase for phosphorylation of S251 (41, 42), are also buried. Therefore, accessibility of S251 and the CKII recognition sites requires a conformational change in the L0 linker, which may occur upon proteolytic cleavage or binding of nucleotide. This sort of rearrangement of the L0 linker upon nucleotide binding has been seen in other ABC proteins (39). Because the L0 linker binds the R region (see below), phosphorylation of the R region could also affect the accessibility of S251. Consequently, phosphorylation of the L0 linker at S251 may fine tune the Ycf1p activity that results from nucleotide binding and hydrolysis, R region phosphorylation, or proteolytic digestion (28, 36). This interplay between different regulatory sites would also be expected to occur in other ABCC proteins, such as MRP1, which is similarly phosphorylated at the L0 linker and R region (43, 44).

### Variable association with the ABC core suggests how TMD0 modulates function in ABC proteins

Although TMD0 helices could not be assigned in the earlier cryo-EM map of MRP1 (22), fitting of the Ycf1p TMD0 model into the MRP1 density map reveals the identity of the TM helices in MRP1 (Fig. S8B). Comparison of the TMD0 structures from Ycf1p, MRP1 (22), and SUR1 (38) shows that the domain adopts the same fold in all three proteins (Fig. 2B, left). The RMSDs between Cα atoms from Ycf1p and MRP1 TMD0, Ycf1p and SUR1 TMD0, and MRP1 and SUR1 TMD0 are 0.80 Å, 0.89 Å, and 0.96 Å, respectively. Despite this similarity, the orientation of TMD0 relative to the ABC core differs dramatically in the three proteins (Fig. 2B, right). The most striking difference is between Ycf1p and SUR1, with a ~145° rotation of TMD0 between the structures (Fig. 2B, bottom right). In SUR1, unlike Ycf1p, residues in TM2 from TMD0 mediate interactions with TM bundle 1 on the extracellular side of the membrane domains (equivalent to the vacuolar lumen side in Ycf1p) and residues in TM2 and TM3 contact the L0 linker on the cytoplasmic side (19, 20). Another consequence of the different orientation of TMD0 is that residues N-terminal to TM1 contact the Kir6.2 in SUR proteins while they contact the ABC core in Ycf1p.

The association of TMD0 with the ABC core also differs between Ycf1p and its homolog MRP1 (Fig. 2B, middle right, and Fig. S8A). While there is no relative rotation of TMD0 between MRP1 and Ycf1p, TMD0 of MRP1 projects away from the ABC core at the extracellular side of the membrane, with distances between backbone atoms in TM4 of TMD0 and TM7 of TM bundle 1 of 16.5 Å in MRP1 versus 9.7 Å in Ycf1p. As a result there are no observed interactions of helical residues in TMD0 with the ABC core on the extracellular side of MRP1, while these interactions occur in Ycf1p (Fig. 2A, top right). However, direct contacts between TMD0 and TM bundle 1 in MRP1 may be formed by loops in TMD0 or the N-terminal sequence of TMD0. Indeed, in the MRP1 map there is a previously-unidentified region of weak density between TMD0 and TMD1 that, from fitting the Ycf1p model of TMD0, appears to correspond to the N-terminal residues of MRP1 TMD0 (Fig. S8B). Thus, both the Ycf1p model and the published MRP1 map suggest that N-terminal residues are important for TMD0/ABC core interactions. Further, the similarity in the orientation of TMD0 relative to TM bundle 1 in Ycf1p and MRP1 suggests that this orientation is preserved among MRP and MRP-like ABCC transporters.

The limited protein-protein interactions between TMD0 and the ABC core in Ycf1p suggest that TMD0 is attached flexibly to the ABC core. It is possible that interactions between TMD0 and the ABC core are altered in different functional states of the transporter, which could affect the orientation and/or dynamics of TMD0. Modest changes in TMD0 orientation are seen for SUR1 TMD0 in KATP channel structures under different nucleotide-bound states (39) and differences in TMD0 density in cryo-EM maps of MRP1 obtained at different points in its catalytic cycle imply that TMD0 dynamics can change (22–24). Another possibility is that TMD0 adopts different orientations in Ycf1p, MRP1, and SUR1 in order to perform various functions in these ABCC proteins. For example, while TMD0 in Ycf1p and a related MRP (MRP2) are required for efficient trafficking of these ABCC proteins (9, 36), SUR1 TMD0 mediates interactions of this ABCC protein with the Kir6.2 pore (19, 20).

### Regulatory (R) region interactions explain its regulatory role

The linker between NBD1 and TMD2 is sometimes referred to as the regulatory (R) region in ABC proteins because its phosphorylation (44, 45) or sumoylation (46) modulates protein activity or expression, respectively. In addition, the R region can serve as an interaction hub for associating proteins (30, 47–50). As seen for intrinsically-disordered regions in other ABC proteins (3, 4), high-resolution density is absent for most of the R region in Ycf1p. However, the map contains high-resolution density for two segments of R region (Fig. 3A, *red*). One of these R region segments consists of residues at the C-terminal end of the R region that lead into the elbow helix of TM bundle 2 (Fig. 3A, left and 3B, left). These Ycf1p residues adopt an extended structure and mediate interactions with the L0 linker. The other well-resolved segment of the R region is located on the peripheral face of NBD1, opposite the ATP binding and NBD dimerization interfaces (Fig. 3A, right and 3B, center and right). This R region segment interacts with residues in NBD1 and in the cytoplasmic extension of TM16. Notably, the R region contacts observed in Ycf1p differs from those observed for other ABCC proteins in the absence of nucleotide and/or substrate (8, 51–53). Interactions of the R region with the peripheral face of NBD1 have only been observed in structures of CFTR where the NBDs are dimerized (54, 55).

**Fig. 3.**
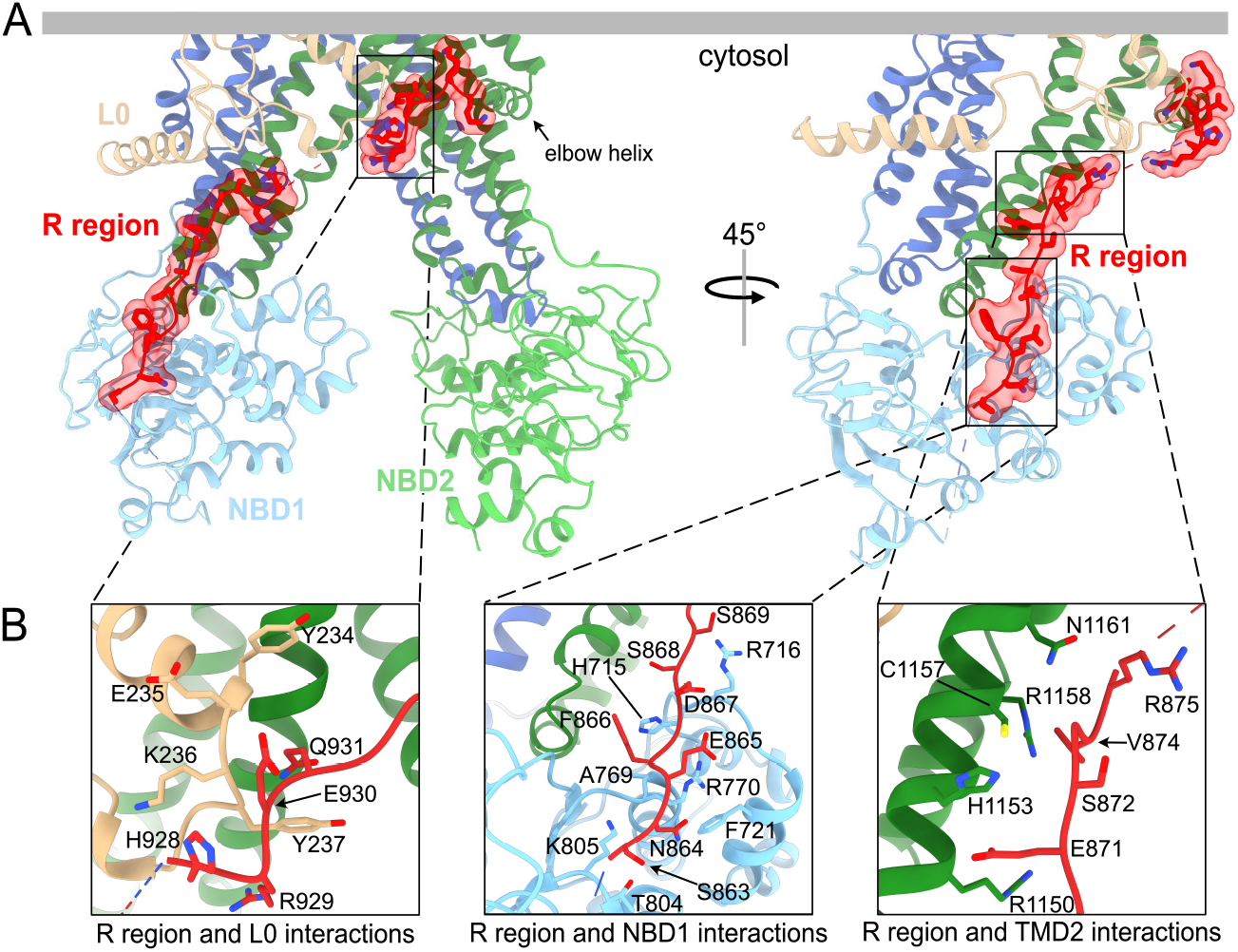
Ycf1p regulatory (R) region interacts with the ABC core and L0 linker. (A) The Ycf1p structure in two orientations with the resolved portions of the R region shown in red. (B) Close up views of the interactions between the R region and the peripheral side of the L0 linker (left), NBD1 (center), and the cytoplasmic extensions in TMD2 (right). Side chains of interacting residues, defined as those within 4 Å of the interacting domain, are shown as sticks.

It is possible that the other portions of the R region are not resolved because of flexibility, but that they still mediate transient interactions with other parts of Ycf1p, such as NBD1, as found with intrinsically-disordered regions of other ABC proteins (30, 48, 56–58). Thus, we conducted tryptophan (Trp) fluorescence quenching studies of isolated NBD1 and NBD1 linked to the R region (NBD1-R) using iodine (I-). Because Trp residues are only found in NBD1, and because I- only quenches Trp residues on the surface of proteins that are surrounded by positively charged residues, these experiments probe changes in the exposure of W701 in Ycf1p NBD1 when the R region is present (59) (Fig. S9A). Notably, the Stern-Volmer constants (KSV) obtained for I- quenching are smaller for NBD1-R than for NBD1 without the R region (Fig. S9B), in both the absence (4.84 ± 0.04 M^−1^ *vs.* 4.59 ± 0.05 M^−1^) and presence of MgATP (3.88 ± 0.04 M^−1^ *vs.* 3.41 ± 0.04 M^−1^), indicating that the surface accessibility of W701 is decreased by the presence of the R region, likely through direct binding of the R region. While this interaction of W701 with the R region is not observed in the Ycf1p model, the fluorescence quenching data suggest that residues outside the two well-resolved R region segments also contact NBD1. Based on the location of W701, we surmise that additional contacts with NBD1 are mediated by residues in the large unresolved E876-E927 segment of the R region, which also contains the two stimulatory phosphorylation sites of Ycf1p. Contact between these residues in the R region and W701 in NBD1 could then be altered upon phosphorylation, as found for phospho-regulatory disordered regions in other ABCC proteins (30, 48, 56–58). Together, the cryo-EM map and fluorescence quenching show the dynamic nature of the R region and its ability to interact with different regions of the ABC protein. These properties enable the R region to exert its known regulatory function on Ycf1p activity.

### Ycf1p structure reveals the molecular basis of human diseases

The atomic-level insight into TMD0 and R region interactions in the Ycf1p model shed light on the underlying basis of disease-causing mutations in human MRPs, such as MRP2 and MRP6 for which this structural information is not available. Mutations in MRP2 (ABCC2) that affect transport of bilirubin glucuronides cause jaundice, choleostisis, and Dubin-Johnson syndrome (60). Mutations in MRP6 (ABCC6) manifest as pseudoxanthoma elasticum (PXE) (61). Disease-causing mutations of MRP2 and MRP6 are found throughout the transporters (Fig.4A; http://www.hgmd.cf.ac.uk/ac/index.php).

The Ycf1p structure shows the locations of residues that are mutated within TMD0 (Fig. 4B, *royal blue*). Several MRP6 missense mutations that cause PXE are located within transmembrane helices in TMD0 and likely affect packing of individual helices in TMD0, destabilizing the domain. PXE-causing mutations in the N-terminal tail and in loop 3 of TMD0 may affect associations of TMD0 with the lumenal side of the ABC core and the L0 linker, respectively. Considering the small number of contacts between TMD0 and other regions of the transporter, even a single mutation would alter how TMD0 interacts with the rest of the ABC transporter. Destabilization of TMD0 and/or altered TMD0/TM helix bundle 1 contacts could have implications for MRP6 trafficking and protein-protein interactions, due to this role for TMD0 in ABC proteins (9, 10, 12–14).

**Fig. 4.**
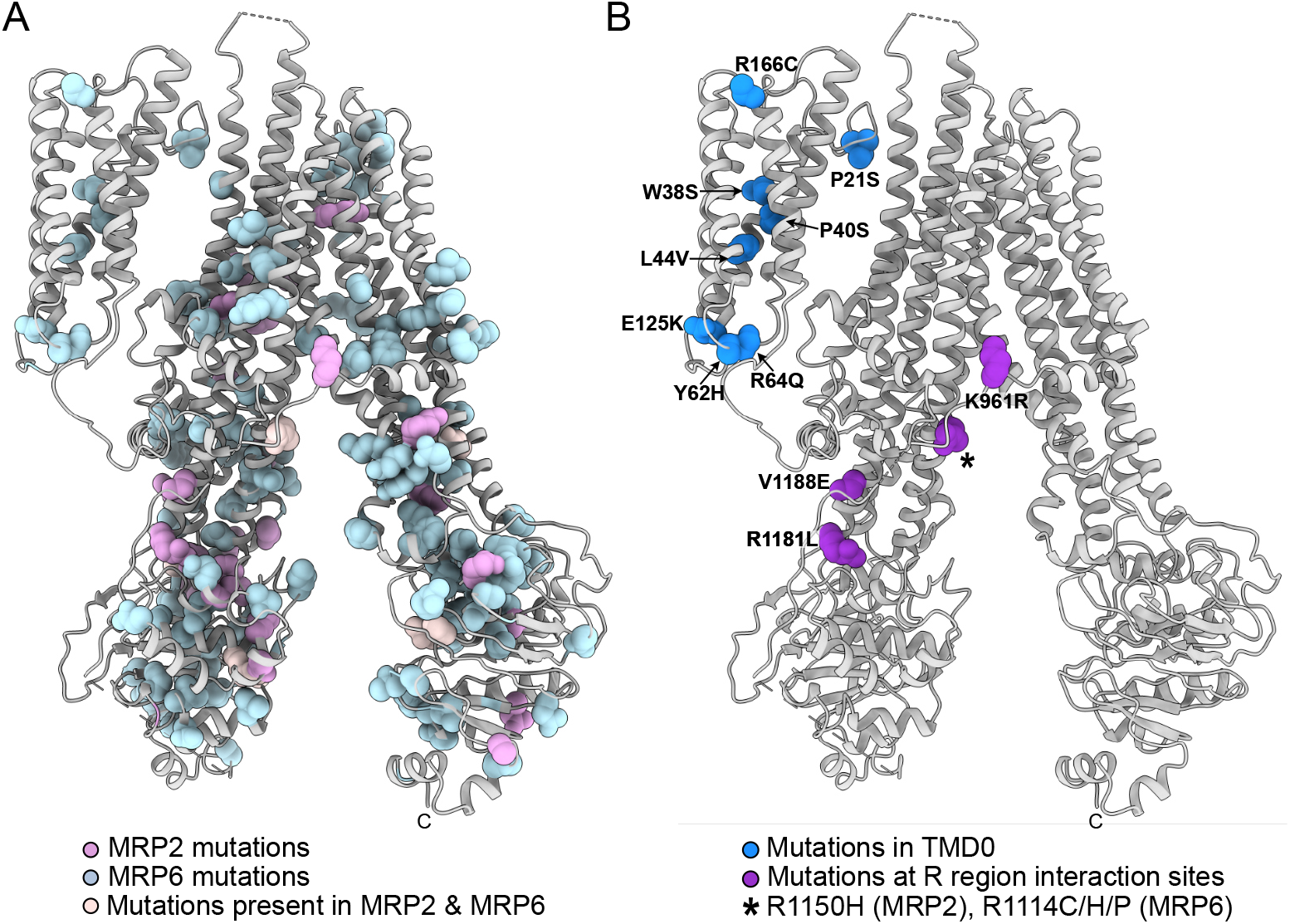
MRP2 and MRP6 disease mutations mapped onto the Ycf1p model. Ycf1p residues that are homologous to MRP2 and MRP6 residues mutated in disease are displayed as spheres and coloured as follows: (A) MRP2 mutations that cause biliary cirrhosis, choleostasis, Dubin-Johnson syndrome, or drug resistance are shown in pink. MRP6 residues altered in in pseudoxanthoma elasticum (PXE) are shown in light blue. Disease-causing mutations of homologous residues in both MRP2 and MRP6 are shown in pale pink (B) Disease-causing mutations of residues in TMD0 are shown in royal blue, while mutations at R region interaction sites are in purple. The specific mutation is listed next to the homologous Ycf1p residue. A black “ * ” indicates a conserved Arg residue that is altered in both MRP2 and MRP6 to cause disease, with the specific mutations indicated.

Other MRP2 and MRP6 mutations likely compromise R region contacts in the transporters (Fig. 4B, *purple*). These mutations are found in the second observed R region segment or in the cytoplasmic extensions of TM16 that contact the R region. Several of these mutations replace Arg residues with hydrophobic residues or introduce charged residues at hydrophobic positions, which would alter interaction of the R region and TMD2. Other mutations substitute Arg residues with other positively-charged amino acids, which subtly alters the charge distribution to affect protein-protein (62) and protein-lipid interactions (63). Cytoplasmic extensions of transmembrane helices in TMD1 and TMD2, and their associated coupling helices (Fig. S1), link ATP binding and hydrolysis at the NBDs to substrate transport. Thus, changes in how the R region contacts the cytoplasmic extensions of TMD1 and TMD2 could alter substrate transport.

As summarized above, the structure of TMD0 and the R region from Ycf1p not only explains the function of this yeast cadmium transporter, but also gives general insight into the ABCC family of proteins. Mapping of MRP mutations onto TMD0 and the R region of Ycf1p structure suggests how these mutations cause disease. The variable interaction between TMD0 and the ABC core shows how this domain can serve different roles in different ABCC proteins while binding of the R region to both the L0 linker and ABC core suggests how its phosphorylation modulates ABCC transporter activity. Together, these insights help explain how nucleotide state and post-translational modifications allow the various component parts of ABCC proteins work together in diverse cellular contexts.

## Materials and Methods

### Construction of the Ycf1p-3×FLAG yeast strain

The *Saccharomyces cerevisiae* strain BJ2168 (MATa, leu2 trp1 ura3-52 prb1-1122 pep4-3 prc1-407 gal2) was further modified by insertion of sequence encoding a C-terminal 3×FLAG tag followed by a *URA3* marker downstream of the *YCF1* gene. The 3×FLAG tag and *URA3* gene were amplified from the pJT1 plasmid (64) and integrated into the yeast chromosome by homologous recombination (65).The sequences of the forward and reverse primers used were 5’-<u>AATCATTGTTCTATTCACT</u>GTGCATGGAGGCTGGTTTGGTCAATGAAAATGACTACA AAGACCATGACGG-3’ and 5’-CCATCCTACGTACCAGATTGTGCGGGACAGGTTTTTATTAGTTTCACAGTTTATAATTGGCCAGTTTTTTTCAAA-3’, respectively, where *YCF1* nucleotides are underlined. The resulting PCR products were used to transform BJ2168 cells, with selection of transformants on minimal media lacking uracil. Successful integration of the sequence for the 3×FLAG tag and *URA3* marker was confirmed by PCR using a primer corresponding to DNA 193 base pairs upstream from the last coding nucleotide of the *YCF1* gene (5’-CGATTCGTACTGCTTTCAAG-3’) and a primer consisting of the reverse and complement of a sequence from within the inserted sequence (5’-GAGCGACCTCATACTATACC-3’). The resulting strain was named ySCB1.

### Purification of Ycf1p

The yeast strain ySCB1 was grown in 11L YPD medium (20 g/L glucose, 20 g/L peptone, and 10 g/L peptone) supplemented with 100 μg/ml of ampicillin, and 0.02% antifoam at 30 °C in a New Brunswick BioFlow fermentor. Yeast was grown for ~22 h to an OD_660_ of 2.0 to 6.0. Cells were harvested by centrifugation at 4,000 g for 15 min at 4 °C and immediately resuspended in 1 ml/g lysis buffer (8 g/L NaCl, 0.2 g/L KCl, 1.44 g/L Na_2_HPO_4_, 0.24 g/L KH_2_PO_4_, 80 g/L sucrose, 20 g/L sorbitol, 20 g/L glucose, 5 mM 6-aminocaproic acid, 5 mM benzamidine, 5 mM EDTA, and 10 mg/L PMSF at pH 7.4). The resuspended cells were lysed with a bead beater (Biospec), cellular debris was removed by centrifugation at 4,000 g for 15 min at 4 °C, and membranes collected by ultracentrifugation (Beckman L-90K) at 152,947 g for 40 min at 4 °C. Membranes were resuspended in 0.5 ml lysis buffer per gram of harvested yeast cells using a Dounce homogenizer and stored at −80 °C prior to protein purification. Frozen membranes were thawed at room temperature and all further purification steps proceeded at 4 °C. Membranes were solubilized with 1 % (w/v) n-dodecyl-β-D-maltopyranoside (DDM, Anatrace) with mixing for ~1 h. Insoluble material was removed by ultracentrifugation at 181,078 g for 70 min (Beckman L-90K). Supernatant containing Ycf1p was filtered with a 0.45 μm syringe filter (Pall) and applied to a 500 μL M2 affinity gel column (Millipore Sigma) pre-equilibrated with DTBS (50 mM Tris-HCl, 150 mM NaCl, 0.04 % (w/v) DDM, pH 8.0). The column was washed with 12 column volumes of DTBS, followed by 8 column buffers of GTBS (50 mM Tris-HCl, 150 mM NaCl, 0.006 % (w/v) glyco-diosgenin (GDN, Anatrace), pH 8.0). The Ycf1p was eluted with 3 column volumes of GTBS containing 150 μg/ml 3×FLAG peptide. An additional 5 column volumes of GTBS without 3×FLAG peptide was applied to wash residual Ycf1p from the column. Fractions containing Ycf1p were concentrated with a 100-kDa MWCO Amicon Ultra centrifugal filter (Millipore Sigma) to a volume ~100 μL, and then applied to a Vivaspin® 500 100-kDa MWCO concentrator and concentrated to ~1.5 mg/ml for ATPase assays and ~5.5 mg/mL for cryo-EM. Protein concentrations were determined by the BCA assay.

ATPase activity of freshly purified Ycf1p in GTBS was measured at 30 °C as described previously (66, 67). Briefly, a reaction mixture was prepared containing 50 mM HEPES, 3 mM MgCl_2_, 0.006% (w/v) GDN, pH 8.0, ~30 μg of Ycf1p, 2 mM ATP, 1 mM PEP, 3.2 units of pyruvate kinase, 8 units of lactate dehydrogenase, and 200 μM NADH. The amount of NADH present in the assay was determined spectrophotometrically at 340 nm from a standard curve in the ATPase assay mixture.

### Cryo-EM and image processing

Nanofabricated holey gold grids were prepared as described previously (68). Grids were glow discharged in air (PELCO easiGLOW) for 2 min immediately before use. A 3 μL sample of the purified Ycf1p at ~4-7 mg/ml was applied in a modified Vitrobot grid preparation robot (FEI) at ~100% relative humidity and 4 °C, and equilibrated for 20 s before blotting both grid sides for 28 s and plunge freezing in a liquid ethane-propane mixture (69). During optimization of cryo-EM grids, specimens were screened at 200 kV with an FEI Tecnai F20 electron microscope equipped with a Gatan K2 Summit camera. High resolution data was collected at 300 kV with a Titan Krios G3 electron microscope (Thermo Fisher Scientific) equipped with a prototype Falcon4 camera. Movies were recorded for 9.6 s with a calibrated pixel size of 1.03 Å, a total exposure of ~45 e^−^/Å^2^, and an exposure rate of 5 e^−^/pixel/s. A dataset of 4,084 movies was collected with EPU software and data collection was monitored with cryoSPARC Live (70). All image processing was performed with cryoSPARC v.2. Patch motion correction and CTF estimation were performed, followed by particle selection by template matching with a circular template and individual particle motion correction. A total of 2,789,383 particle images were used for 2D classification. Subsequent *ab initio* structure determination, 3D classification, and non-uniform refinement led to a final map at 3.2 Å resolution from 124,864 particle images.

### Model Building and Validation

An initial model of Ycf1p based on the known structure of bovine MRP1 (5UJ9, (22)) was generated with Phenix (71, 72). Coot (73) was used for manual atomic model building, with real-space refinement in Phenix (71, 72). The TMD0 domain, the observed segments of the R region, and several intracellular and extracellular loops were built *de novo*. The structure of TMD0 was determined by building transmembrane helices of TM1, TM4, and TM5 first, followed by the N-terminal tail, TM2 and TM3, and the TMD0 loops. Loop 5, which connects TMD0 to the L0 linker and for which only weak density is observed, was built into the model based on ideal backbone Ramachandran angles. The first observed segment of the R region was built based on the side chains of F866, E8871, S872, and V874. Homology models of NBD1 and NBD2 were generated in Modeller (74) based on a structure-based sequence alignment of NBD1 or NBD2, respectively, from ABCC proteins. The homology model of NBD1 was generated using the structures of NBD1 from MRP1 (PDB: 2CBZ (75)), ABCC6 (PDB: 6NL0), and hamster SUR1 (PDB: 6JB1 (76)). The homology model of Ycf1p NBD2 was generated using the structures of NBD2 from MRP1 (PDB ID: 6BHU (23)) and ABCC6 (PDB: 6BZR). The NBD1 and NBD2 homology models were fit into the low-density NBD regions that possessed defined secondary structure. Phospholipids were added and refined with Coot. Data collection, image processing, and model statistics are in Table S1.

All figures were generated with UCSF ChimeraX (77) and map-in-model fits with USCF Chimera (78). Structure comparisons were done with the ChimeraX matchmaker tool using the default Needleman-Wunsch BLOSUM-62 matrix. Root-mean squared (RMSD) values were determined comparing Cα positions using the Match > Align tool UCSF Chimera. Electrostatic surface representation of NBD1-R was generated in MolMol (79).

### NBD1 and NBD1-R Expression and Purification

Ycf1p NBD1 (D610-N859) and NBD1-R (D610-E930) were expressed and purified as described previously (58, 59, 80). Briefly, Ycf1p NBD1 and NBD1-R were expressed as N-terminal His6-SUMO fusion proteins in *E. coli* LOBSTR-BL21 (DE3) RIL (Kerafast Inc) using a pET26b-derived expression vector (81). Cell cultures were grown in 1 L of 97.5% M9 and 2.5% LB media at 37 °C to an OD_600_ of 0.4 to 0.5, at which point the temperature was gradually lowered so that the OD_600_ was 0.8 when the temperature reached 18 °C. Cell cultures were incubated at 18 °C for 30 min and gene expression was subsequently induced with 0.75 mM isopropyl β-D-thiogalactoside (IPTG). Cells were harvested by centrifugation 18 to 20 h post-induction and stored at −20 °C until purification.

All purification steps were conducted at 4 °C. Cell pellets were incubated on ice for 30 min and then resuspended in 15 ml of lysis buffer per 1 L of harvested cells (20 mM Tris-HCl, 10% glycerol, 150 mM NaCl, 5 mM imidazole, 100 mM arginine, 2 mM β-mecraptoethanol, 15 mM MgCl_2_, 15 mM ATP, 0.2% (v/v) Triton X-100, 1 mg/ml lysozyme, 2 mg/ml DCA, 5 mM benzamidine, 5 mM ε-amino n-caproic acid, 1 mM PMSF, pH 7.6). A trace amount of DNase I was added, and the solution was incubated for 15 min prior to sonication (Misoniz Microson Ultrasonic Cell Disruptor). The soluble and insoluble fractions were separated by centrifugation at 17,000 g for 30 min. The lysate was then filtered using a 0.45 μM Acrodisic syringe filter (Pall) and applied to a 5 mL Ni^2+^-NTA affinity column (GE Healthcare) equilibrated with 20 mM Tris-HCl, 10% (v/v) glycerol, 150 mM NaCl, 5 mM imidazole, pH 7.6. The column was then washed with 5 column volumes of the equilibration buffer that also contained 5 mM MgCl_2_ and 5 mM ATP. The bound His_6_-SUMO-NBD1 or His_6_-SUMO-NBD1-R fusion proteins were eluted with 5 column volumes of elution buffer (20 mM Tris-HCl, 10% (v/v) glycerol, 150 mM NaCl, 500 mM imidazole, 15 mM MgCl_2_, 15 mM ATP, 1 mM benzamidine, 1 mM ε-amino n-caproic acid). The elution fractions were immediately diluted 3-fold with elution buffer lacking imidazole and dialyzed overnight against 20 mM Tris, 10% (v/v) glycerol, 150 mM NaCl, 2 mM β-mercaptoethanol, 2 mM MgCl_2_, 2 mM ATP, 1 mM benzamidine, 1 mM ε-amino n-caproic acid, pH 7.6. His_6_-Ulp1 protease (~1.8 mg) was added to the dialysis bag in order to remove the His_6_-SUMO tag from NBD1 (or NBD1-R). The resulting solution, containing His_6_-SUMO and NBD1 (or NBD1-R) proteins, was loaded onto a Co^2+^-agarose affinity column (Thermo Scientific), which was pre-equilibrated in 20 mM Tris, 10% (v/v) glycerol, 150 mM NaCl, 10 mM imidazole, 10 mM MgCl_2_, 10 mM ATP, pH 7.6, to isolate NBD1 from the His_6_-SUMO tag. Fractions containing Ycf1p NBD1 or NBD1-R were applied to a size exclusion column (Superdex 75, GE Healthcare) in 20 mM Na_3_PO_4_, 150 mM NaCl, 2 % (v/v) glycerol, 2 mM DTT, 2 mM MgCl_2_, 2 mM ATP, 100 μM benzamidine, 100μM ε-amino n-caproic acid, pH 7.3 to purify the protein to homogeneity. All protein concentrations were determined by the Bradford protein assay (82).

### Iodide Trp Fluorescence Quenching Experiments

Fluorescence quenching experiments were performed on 2 μM Ycf1p NBD1 or NBD1-R. Purified Ycf1p NBD1 and NBD1-R were exchanged into the fluorescence buffer (20 mM Na_3_PO_4_, 2% glycerol, 150 mM NaCl, pH 7.3) by size exclusion chromatography (Superdex75, GE Healthcare). MgATP-bound and apo NBD1 and NBD1-R were generated by adding 2 mM MgATP or 2 mM EDTA, respectively, to the protein eluents. Fresh reductant (2 mM DTT) was added to quenching solutions. Fluorescence studies were conducted on a Fluoromax-4 fluorimeter (Horiba Scientific) equipped with a Peltier unit for temperature control. Trp emission spectra (310 nm – 450 nm) were recorded at 15 °C with an excitation wavelength of 298 nm, an excitation slit width of 2 nm, and an emission slit width of 5 nm.

Iodine (I-) quenching experiments were conducted by successive additions of 0.8 M KI dissolved in the fluorescence buffer. The quenching solution was prepared fresh and contained 0.2 mM Na_2_SO_3_ to prevent formation of I_2_ and I_3_^−^(83). A parallel titration using 0.8 M KCl was conducted to account for sample dilution and changes in ionic strength during the titration. The initial fluorescence (F_O_) at each titration point was divided by the fluorescence (F) to give (F_O_/F)_KI_ or (F_O_/F)_KCl_. The change in fluorescence, (F_O_/F)_KI_/(F_O_/F)_KCl_, was plotted against the concentration of I^−^ to determine the K_SV_ constant (84).

## Acknowledgements

The authors thank Stephanie Bueler with assistance in generating the ySCB1 yeast strain, and Thamiya Vasanthakumar and Zev Ripstein for helpful discussions regarding ATPase assays and computation, respectively. SCB was supported by a Canadian Graduate Scholarship-Master’s Program scholarship from the Natural Sciences and Engineering Council (NSERC) and the Sarah Cusick Gallop and William George Gallop Memorial Scholarship. JLR was supported by the Canada Research Chairs program. The research was supported by NSERC grants RGPIN-2015-05372 (VK) and RGPIN-2020-05835 (VK), Heart and Stroke Foundation of Canada grant G-18-0022076 (VK), and Canadian Institutes of Health Research grant PJT162186 (JLR). Cryo-EM data were collected at the Toronto High Resolution High Throughput cryo-EM facility, supported by the Canada Foundation for Innovation and Ontario Research Fund.

## Statements of Contributions

VK conceived of the project and designed experiments with JLR and SCB. SCB created the yeast strain and protein expression plasmids, purified protein, performed ATPase assays and fluorescence studies, processed cryo-EM images, and constructed the Ycf1p atomic model. SB prepared cryo-EM specimens and collected images. Figures were prepared by SCB and the manuscript was written by SCB, VK, and JLR.

## Notes

### Competing Interest Statement

The authors have declared no competing interest.

