## Supplemental Files for "Structure of Ycf1p reveals the transmembrane domain TMD0 and the regulatory R region of ABCC transporters"

This PDF file includes:
Figs. S1 to S9
Table S1
References for supplementary information

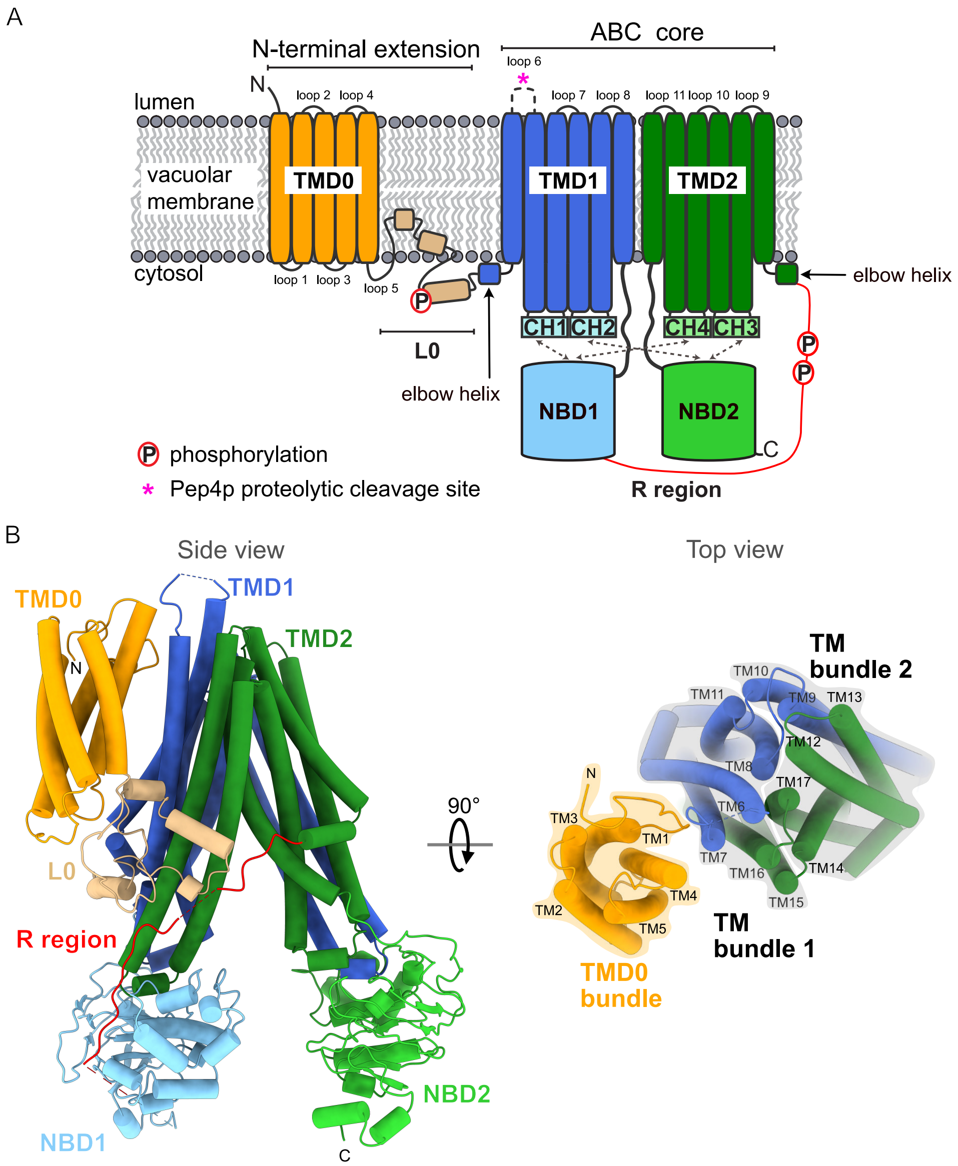


**Fig. S1. Domain architecture and post-translational modifications in Ycf1p.** (A) Cartoon representation of Ycf1p in the vacuolar membrane showing the two transmembrane domains (TMD1 in dark blue and TMD2 in dark green), two nucleotide binding domains (NBD1 in light blue and NBD2 in light green), and regulatory (R) region (red) that comprise the ABC core. Coupling helices (CH1, CH2, CH3, CH4) that connect the cytoplasmic extensions of the transmembrane (TM) helices in TMD1 and TMD2 are also shown, with CH1 and CH2 in pale blue, and CH3 and CH4 in pale green. The elbow helices of TMD1 and TMD2 are shown in blue and green, respectively. The N-terminal extension of Ycf1p is formed by TMD0 (orange) and L0 linker (tan). Loops, either in the lumen or cytosol, are labeled sequentially. Loop 6, which is proteolytically cleaved, is depicted as a dashed curve with a pink asterisk highlighting the proteolysis site. Phosphorylation sites in the L0 linker (S251) and R region (S908 and T911) are depicted with a red-encircled “ P ”. (B) Schematic diagram of the Ycf1p cryo-EM structure with cylinders representing α-helices and arrows representing β-strands. The top view, which is from the lumenal side, highlights the composition and orientation of the TMD0 bundle (TM helices 1-5), TM bundle 1 (TM helices 6,7,8,11,15,16), and TM bundle 2 (TM helices 9,10,12,13,14,17).

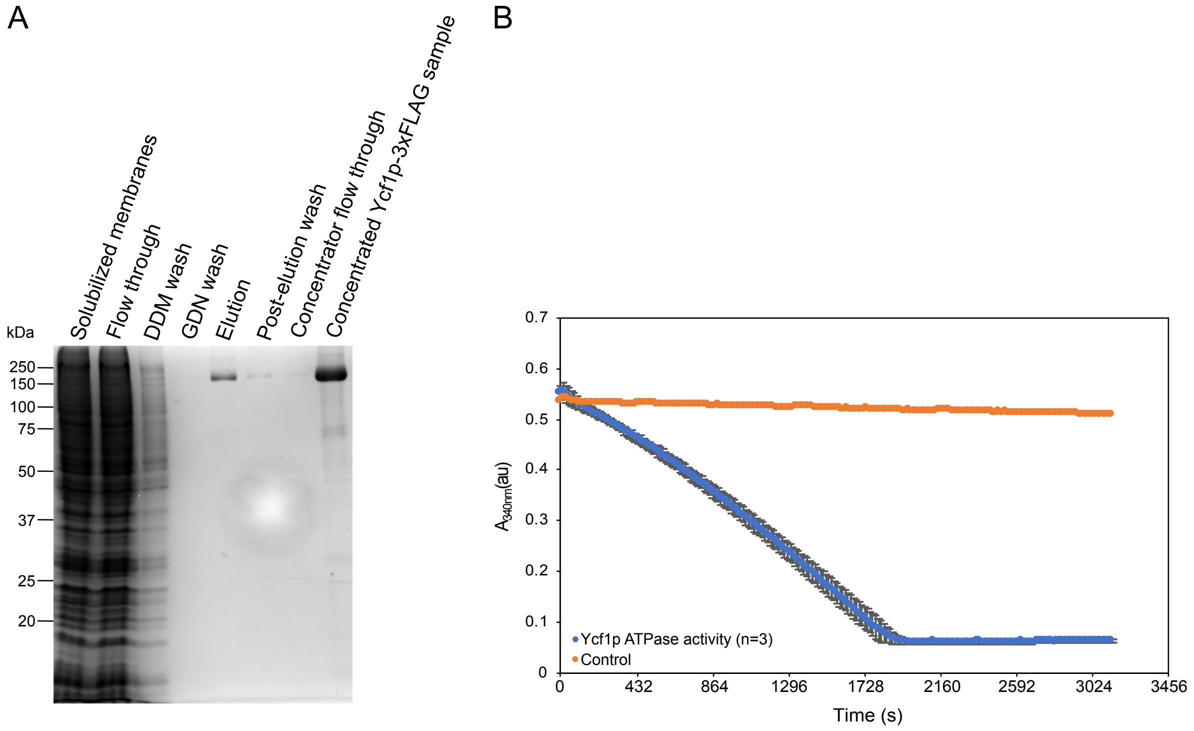


**Fig. S2. Ycf1p sample preparation.** (A) 12% SDS-PAGE gel of a purification of Ycf1p-3×FLAG sample (171 kDa). (B) ATPase activity of Ycf1p. Blue circles indicate the mean value with error bars showing ±s.d. (n=3 assays) for absorbance at 340 nm, which follows oxidation of NADH in an enzyme-coupled assay.

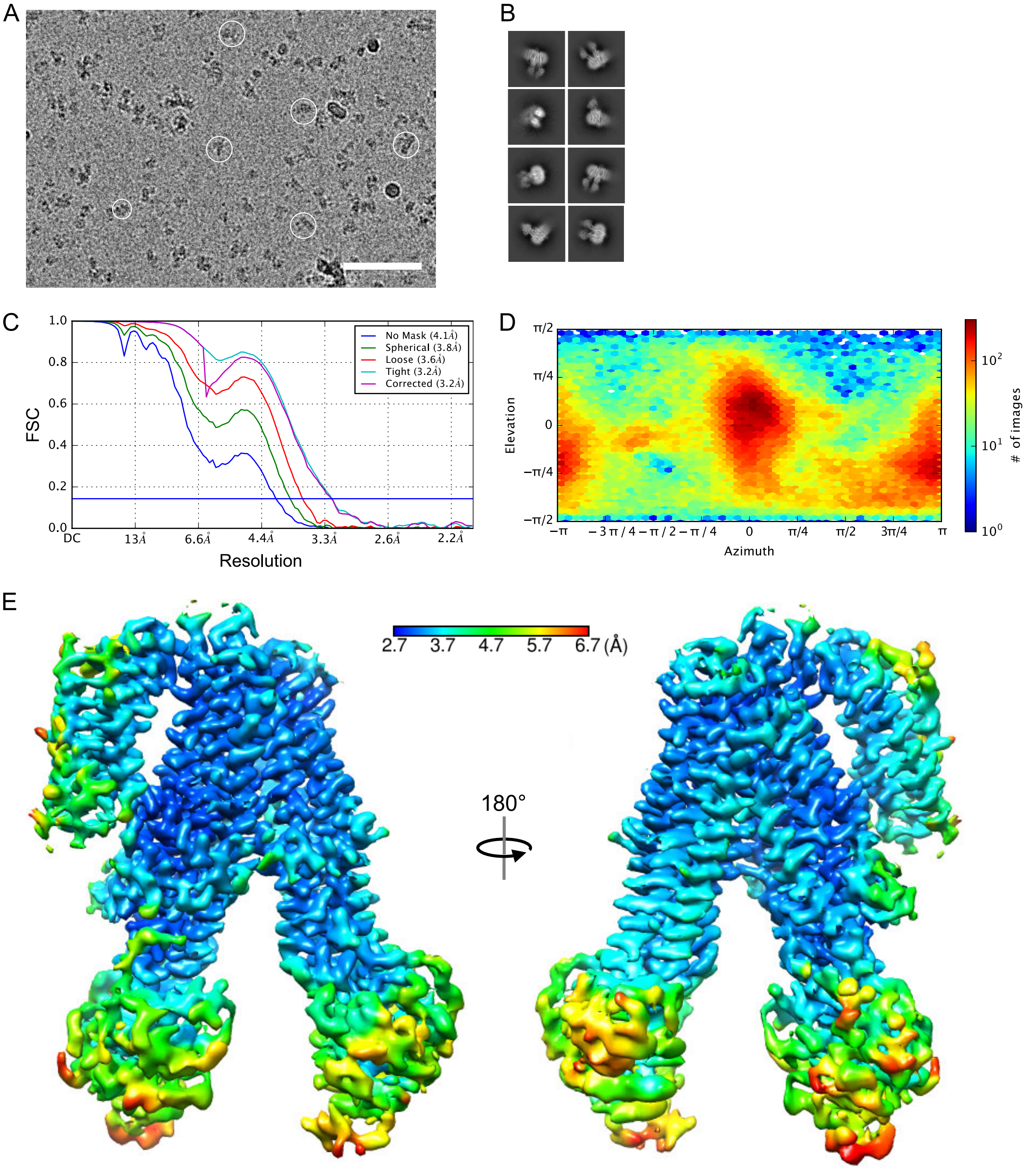


**Fig. S3. Cryo-EM image analysis.** (A) Section of a representative micrograph with example Ycf1p particles circled in white. Scale bar, 500 Å. (B) Example 2D class average images of Ycf1p. (C) Fourier Shell Correlation (FSC) curve for the Ycf1p map. (D) Euler angle distribution for particle images contributing to the map. (E) Local resolution estimated for the Ycf1p map.

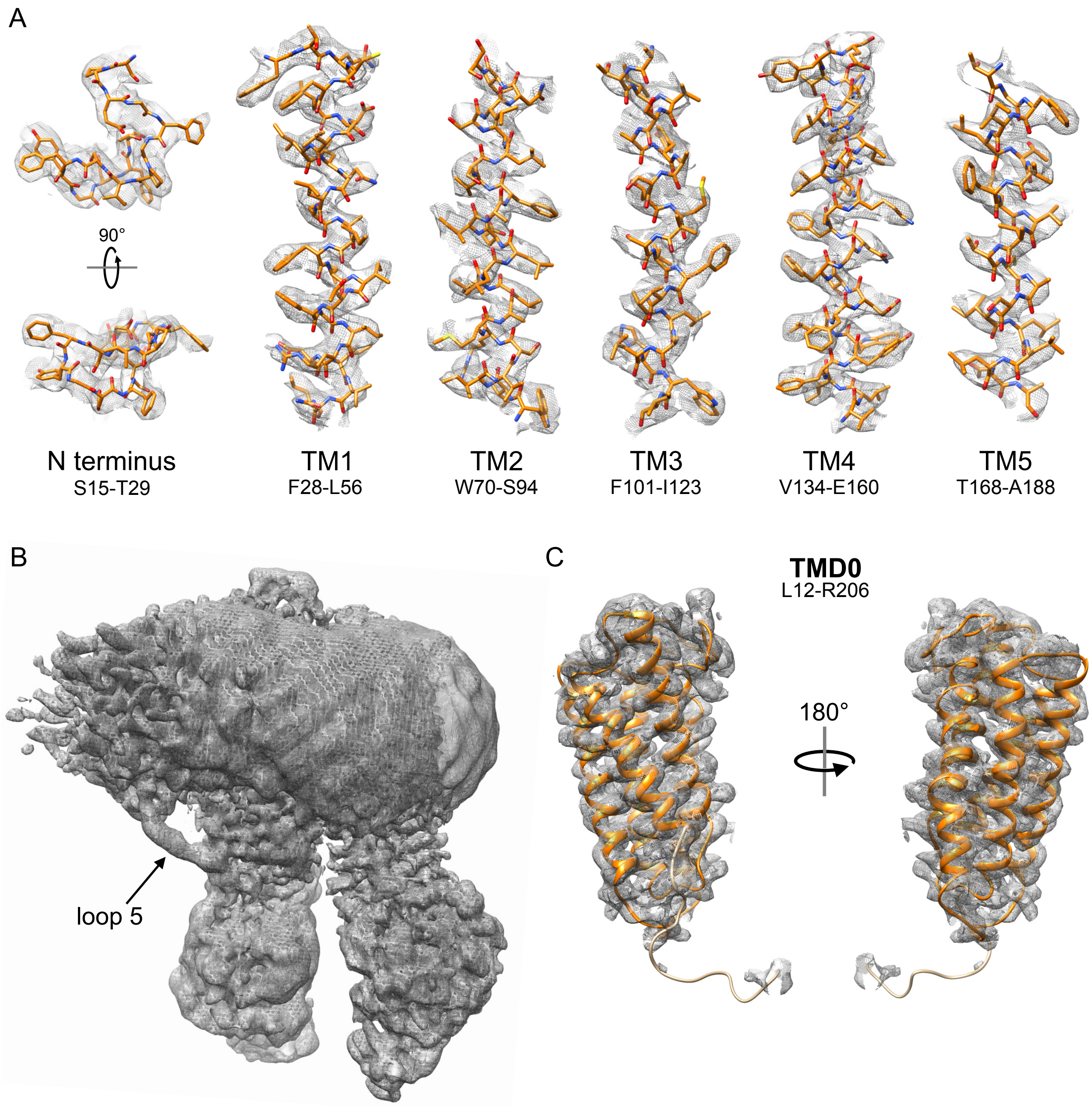


**Fig. S4. Atomic model of TMD0.** (A) Fitting of the lumenal N-terminal tail and individual TMD0 helices (TM1 to TM5) into the cryo-EM density. (B) Low-resolution density corresponding to cytosolic loop 5 (L186-R206), which connects TMD0 and the membrane embedded region of the L0 linker, is seen consistently when maps of Ycf1p are rendered at a low-density cutoff. (C) Model of TMD0 fit into experimental cryo-EM density map.

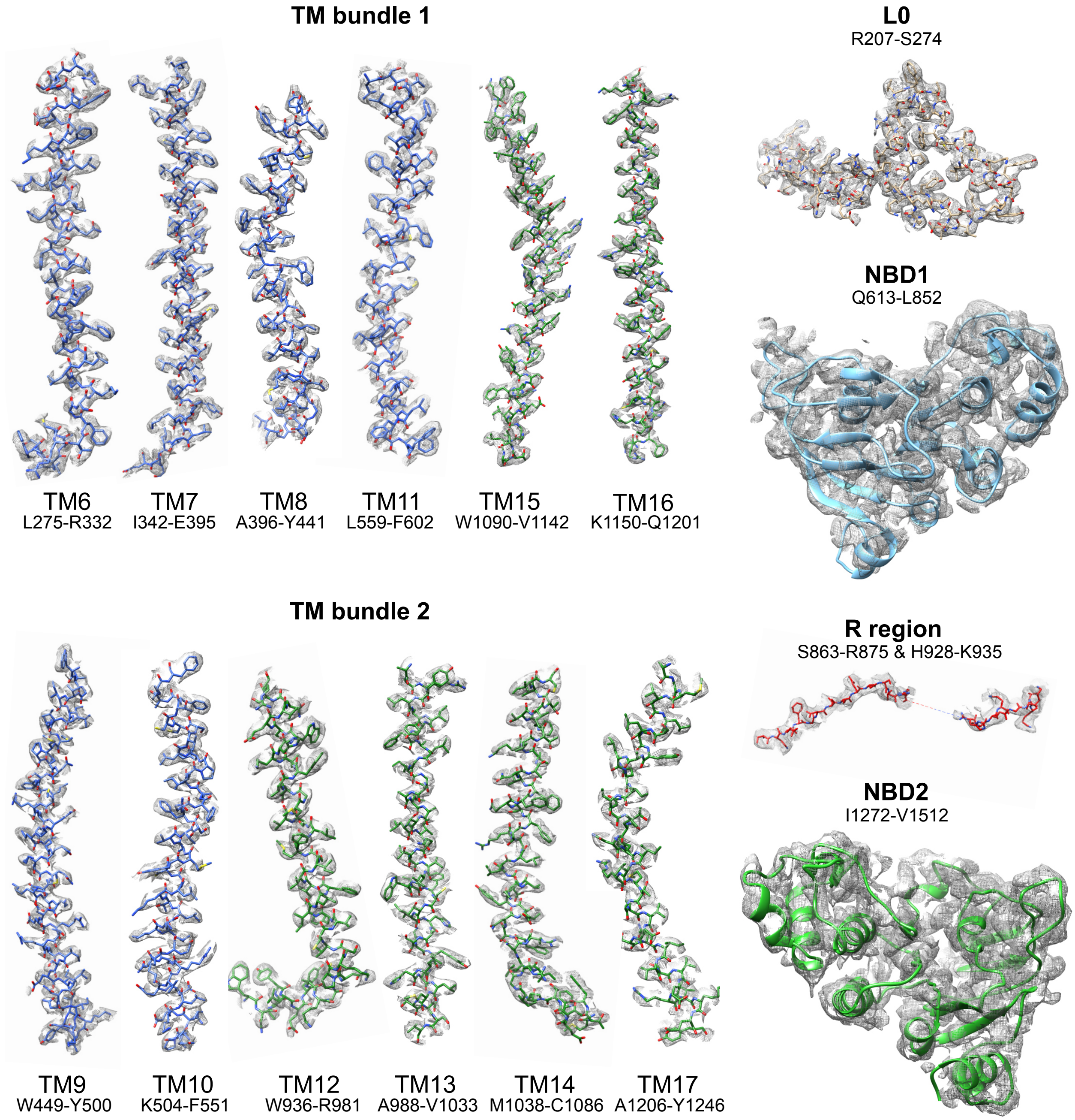


**Fig. S5. Representative model-in-map fit for the L0 linker and ABC core of Ycf1p.** Examples of the atomic model of Ycf1p fit into the experimental cryo- EM map are shown for individual transmembrane helices in TM bundles 1 and 2, the L0 linker, two segments of the R region, and NBD1 and NBD2.

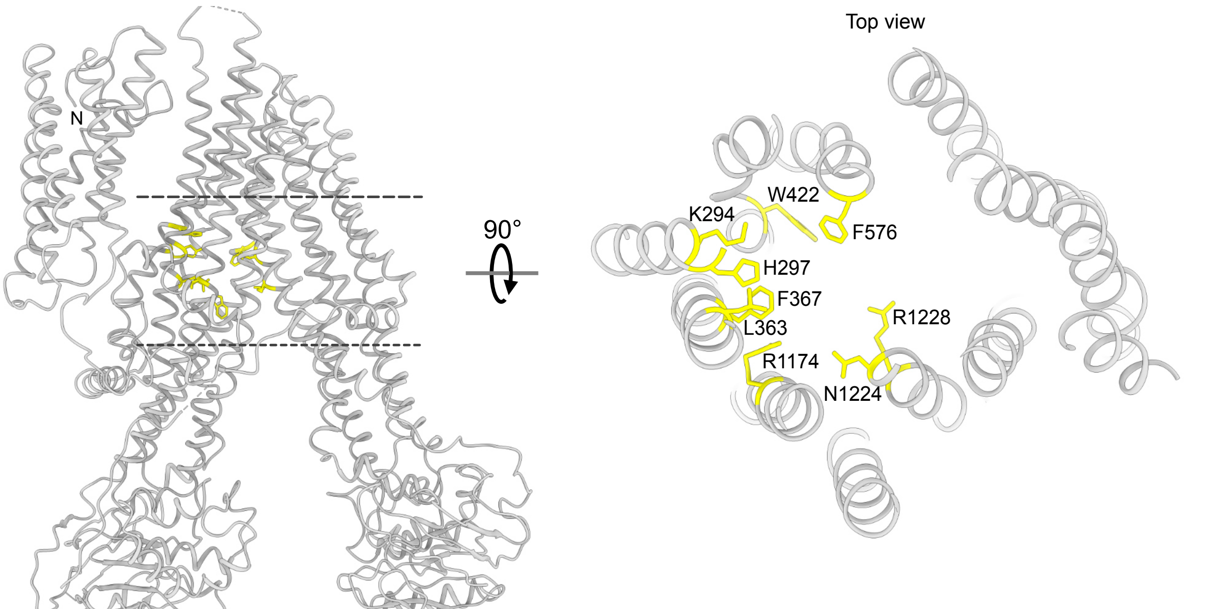


**Ycf1p (bovine MRP1) GSH-binding residues**

| | | |
|---|---|---|
| K294 (K332) | F367 (F385) | R1174 (R1196) |
| H297 (H335) | W422 (Y440) | N1224 (N1244) |
| L363 (L381) | F576 (F594) | R1228 (R1248) |

**Fig. S6. Homologous bovine MRP1 GSH binding residues in Ycf1p.** The identity of potential GSH-binding residues in Ycf1p, shown in yellow, are based on the structure of bovine MRP1 bound to substrate (1). Ycf1p residues that form the putative GSH-binding site are summarized in the table, with the homologous GSH-binding residues in bovine MRP1 indicated in brackets.

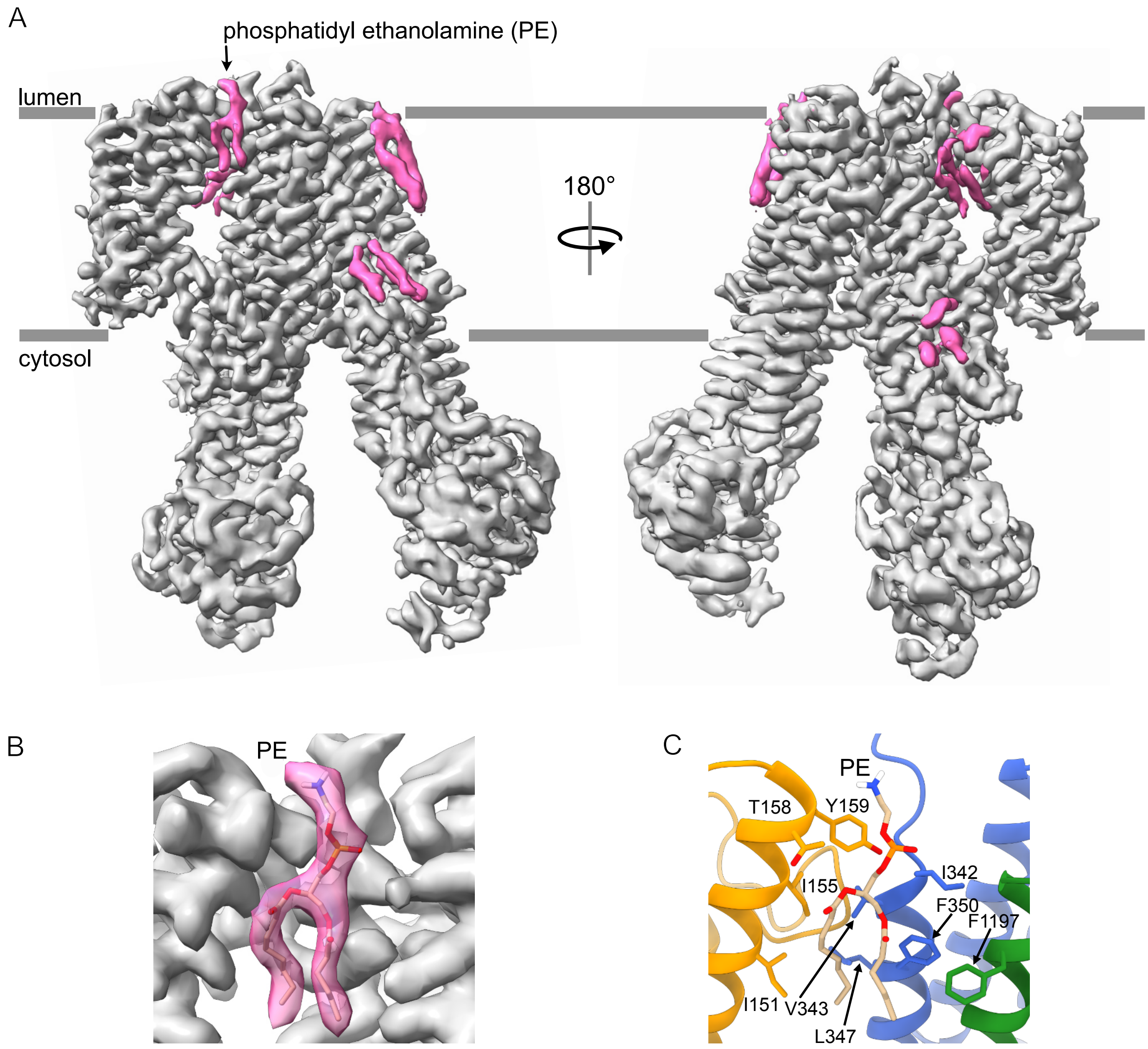


**Fig. S7. Lipid densities in Ycf1p transmembrane region.** (A) Cryo-EM map of Ycf1p with density corresponding to the protein in grey and density corresponding to lipid molecules in pink. The prominent phospholipid density at the TMD0/ABC core interface at lumenal side membrane is modeled as phosphatidylethanolamine. (B) Fit of the phosphatidylethanolamine into the corresponding phospholipid density with the lipid tail truncated to fit the observed density. (C) Interactions of phosphatidylethanolamine with residues in TM4 (I151, T157, Y159), TM7 (I342, V343, F346, L347), and TM15 (F1197). The lipid is shown in tan, TMD0 is shown in orange, TMD1 is shown in blue, and TMD2 is shown in green.

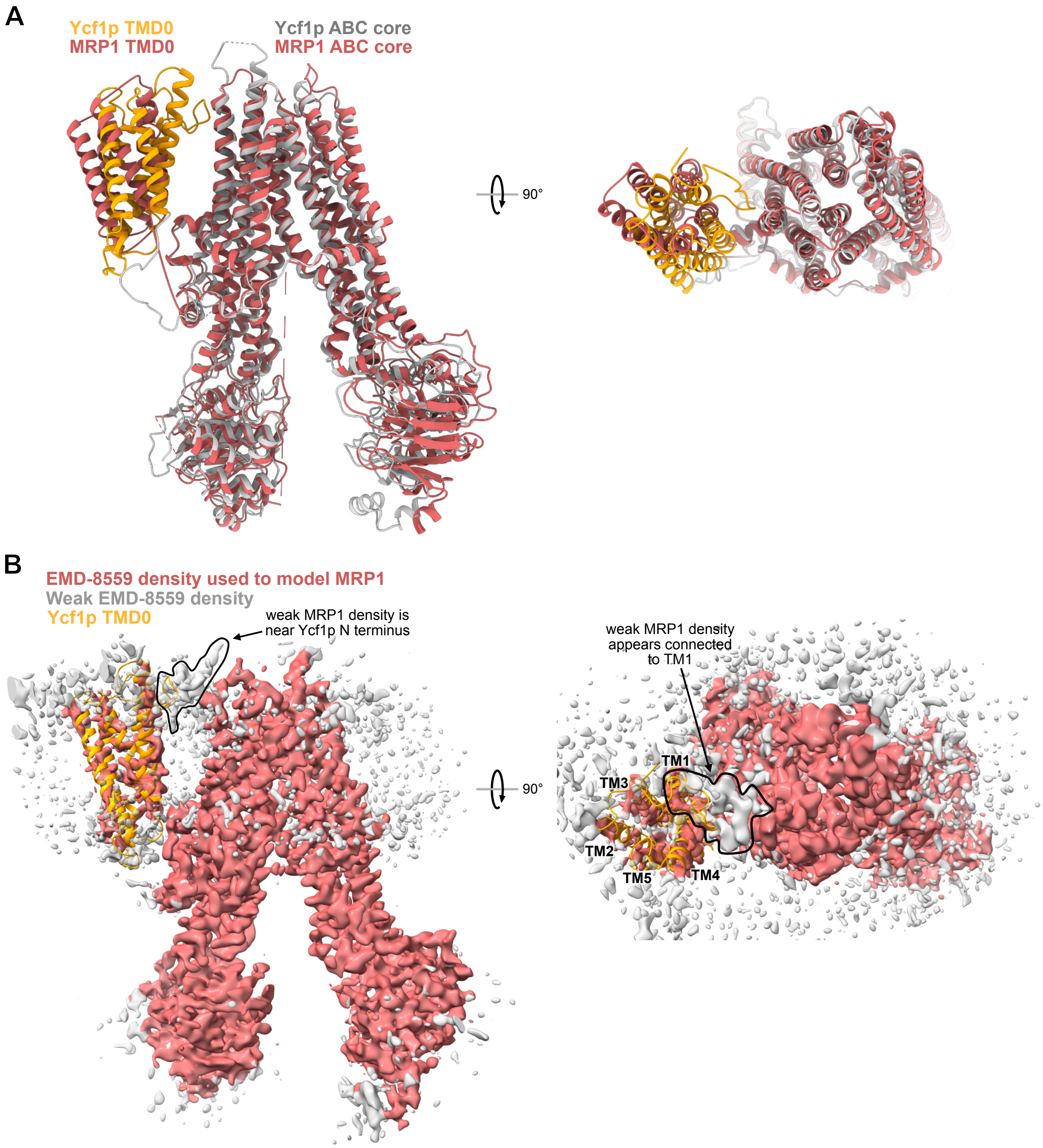


**Fig. S8. Assignment of bovine MRP1 TMD0 helices and the unknown density at the interface between TMD0 and the ABC core extracellular interface based on the Ycf1p structure.** (A) Overlay of the ABC core and L0 linker of Ycf1p and bovine MRP1 (R.M.S.D. 1.88 Å), showing that the TMD0 helix bundle of MRP1 projects away from the ABC core. The MRP1 structure is shown in red while Ycf1p TMD0 is in orange and both the Ycf1p L0 linker and ABC core are in grey. (B) The cryo-EM model of Ycf1p TMD0 (orange) can be fit into the MRP1 density (grey) revealing the topology of MRP1 TMD0. Notably, the region of weak density in the MRP1 map at the extracellular side of the membrane can be tentatively assigned to the N-terminal tail of MRP1.

A

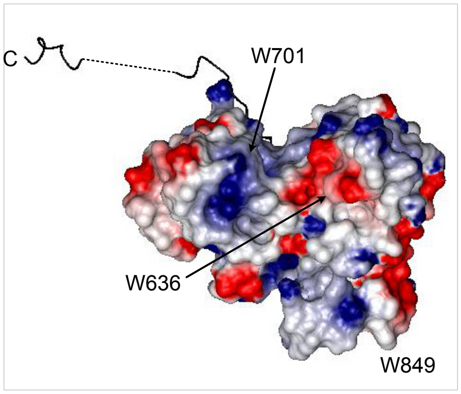


blue = positively charged residues
red = negatively charged residues
white = neutral residues

B

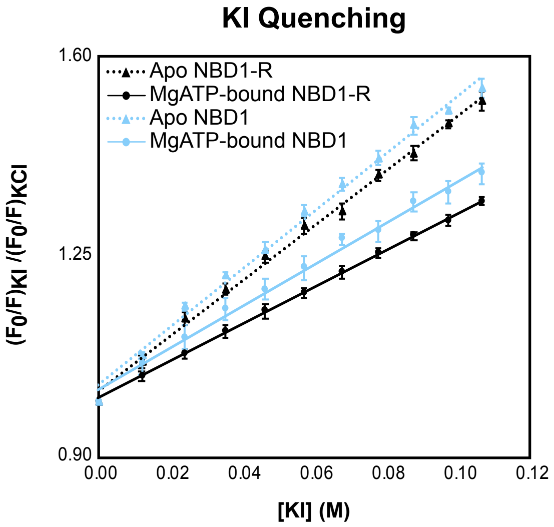


| | NBD1 | | NBD1–R | |
|---|---|---|---|---|
| | Apo | MgATP-bound | Apo | MgATP-bound |
| $K_{sv}$ ($M^{-1}$) KI | 4.84 ± 0.04 (n=3) | 3.88 ± 0.04 (n=3) | 4.59 ± 0.05 (n=3) | 3.41 ± 0.04 (n=3) |

**Fig. S9. Fluorescence quenching for NBD1 with and without the R region.** (A) Surface representation of NBD1 from the Ycf1p atomic model, with blue and red representing positive and negative electrostatic potential, respectively, and white for neutral residues. The three Trp residues in NBD1 are labeled. The backbone traces of the observed segments R region are shown as solid black coils while a dotted black line denotes the unresolved R region segment. (B) Stern-Volmer plot for I- quenching of NBD1 (blue) and NBD1-R (black) in apo (dotted lines) and MgATP-bound (solid lines) states. The table reports the average Stern-Volmer constants ($K_{SV}$) ± s.d. (n=3 independent measurements).

**Table S1. Summary of data collection, image processing, and model statistics**

| **Data Collection** | **EMD-XXXX** |
|---|---|
| Electron microscope | Titan Krios G3 |
| Camera | Falcon4 |
| Voltage (kV) | 300 |
| Pixel size (Å) | 1.03 |
| Total exposure (e-/Å$^2$) | 45 |
| Exposure rate (e-/pixels/s) | 5 |
| Number of frames | 30 |
| **Image Processing** | **EMD-XXXX** |
| Particle collection and selection software | cryoSPARC Live |
| 3D map classification and refinement software | cryoSPARC v2 |
| Motion correction software | cryoSPARC v2 |
| CTF estimation software | cryoSPARC v2 |
| Number of movies used | 4,084 |
| Initial particle images selected (no.) | 2,789,383 |
| Particle images contributing to final map (no.) | 124,864 |
| Symmetry | C1 |
| Global map resolution (Å) | 3.2 |
| **Model Building** | **PDB-YYYY** |
| Modelling and refinement software | *Coot, Phenix* |
| Number of residues built | 1428 |
| RMS bond length (Å) | 0.005 |
| RMS bond angle (º) | 0.681 |
| Ramachadran outliers (%) | 0.0 |
| Ramachadran allowed (%) | 6.9 |
| Ramachadran favoured (%) | 93.1 |
| Rotamer outliers (%) | 0.0 |
| All-atom clashscore | 14.5 |
| MolProbability score | 2.11 |
| EMRinger score | 2.09 |
